# Does Human-Like Contextual Object Recognition Emerge from Language Supervision and Language-Guided Inference?

**DOI:** 10.1101/2025.07.24.666375

**Authors:** Karim Rajaei, Radoslaw Martin Cichy, Hamid Soltanian-Zadeh

## Abstract

Human vision is an active, context-sensitive process that interprets objects in relation to their surroundings. While behavioral research has long shown that scene context facilitates object recognition, the underlying computational mechanisms—and the extent to which artificial vision models replicate this ability—remain unclear. Here, we addressed this gap by combining human behavioral experiments with computational modeling to investigate how structured scene context influences object recognition. Using a novel 3D simulation framework, we embedded target objects into indoor scenes, and manipulated contextual coherence between objects and scenes by using either intact scenes or their phase-scrambled versions. Humans showed a robust object recognition advantage in coherent scenes, particularly under challenging conditions such as occlusion, crowding, or non-canonical viewpoints. Conventional vision models—including convolutional neural networks (CNNs) and vision transformers (ViTs)—failed to replicate this effect. In contrast, vision-language models (VLMs), particularly those using ViT architectures and trained with language supervision (e.g., CLIP), approached human-like accuracy.

This shows that semantically rich and category-structured representations are required for modelling context sensitivity. Notably, context sensitive behavior was closest to humans in VLMs when using language-guided inference at test time. This suggests that how a model accesses its representations during inference is relevant for enabling context-sensitive behavior. Together, this work offers steps towards a computational account of contextual facilitation of objects by scenes, and highlights zero-shot inference as an interesting alignment metric when benchmarking artificial and biological vision.

## Introduction

Imagine being in a crowded living room: you see furniture arranged across the space, a family interacting, everyday objects scattered on tables and shelves. This scene, being visually and semantically rich, is open to countless interpretations. Yet, humans effortlessly make sense of the scene by drawing on both visual input and prior visual knowledge (Gilbert & Li, 2013; Kreiman & Serre, 2020; Malcolm et al., 2016). This ability highlights a central insight from cognitive neuroscience about human vision: it is as much about constructing meaning top-down using prior conceptual knowledge (Collins & Olson, 2014; Gauthier et al., 2003; Lupyan et al., 2010) as it is about bottom-up sensory processing.

A key facet of meaning construction using prior knowledge is making use of contextual regularities to constrain perceptual interpretation (Bar, 2004; Peelen et al., 2024). This is based on the statistical fact that objects rarely appear in isolation, but rather in context that obeys predictable patterns (Peelen et al., 2024; Võ, 2021). For example, plates and utensils co-occur in kitchens, books are found on shelves. Extensive research indicates that context facilitates object perception behavior (Biederman, 1972; Davenport & Potter, 2004; Markov & Võ, 2025; Munneke et al., 2013; Oliva & Torralba, 2007), and neuroscientific findings show that the brain exploits these regularities, integrating local features with global scene context to guide perceptual inference (Aminoff & Tarr, 2015; Bar, 2004; Brandman & Peelen, 2017; Kaiser et al., 2019; Wischnewski & Peelen, 2021).

However, the computational mechanisms behind this integration remain unclear (Peelen et al., 2024). In particular, unlike humans, current artificial vision models often struggle to recognize objects correctly when they appear in unusual configurations or occluded settings (Jacob et al., 2021; Peters & Kriegeskorte, 2021). This gap motivates our central question: under what conditions do artificial models approximate human-like contextual sensitivity?

Previous research has shown that deep neural networks (DNNs) trained on large labeled visual datasets like ImageNet achieve impressive object recognition performance and predict both human brain activity and behavior well (Cichy et al., 2016; Khaligh-Razavi & Kriegeskorte, 2014; Rajaei et al., 2019; Yamins et al., 2014; Zhuang et al., 2022). However, these models often fall short in real-world settings where object recognition depends not only on local features but also on the broader scene context (Jacob et al., 2021; Peters & Kriegeskorte, 2021). We hypothesize that this is to because such purely vision-guided models do not capture and use the contextual information when needed to robustly solve visual object recognition in context. In contrast, vision-language models (VLMs) such as CLIP integrate visual features with language descriptions, allowing them to learn semantically structured, multimodal representations (Radford et al., 2021) that capture complex visuo-semantic information (Doerig et al., 2024; Wang et al., 2023).

Importantly, they also support language-guided inference via zero-shot classification, where images are matched directly to category-level text prompts. We hypothesize that this combination of semantic supervision and flexible inference enables VLMs to apply their knowledge in an adaptive, context-sensitive manner that is unavailable for vision-guided DNNs.

In this study, we determine how scene context affects object recognition in humans and benchmark current DNNs against human behavior. For this, we first conducted a psychophysics experiment in humans, establishing that context impacts object recognition specifically when object viewing conditions are challenging. We then evaluated a wide range of models—from purely vision-guided to language-guided models—to assess whether and how contextual facilitation emerges.

Our findings reveal that VLMs applying knowledge via language-guided inference are incompletely, yet arguably more closely approximating human-like use of context than other models and inference procedures. This highlights the combined importance of semantically rich visual representations and inference procedure in modelling human visual context sensitivity, suggesting relevant constraints for benchmarking.

## Results

### Contextual Structure Enhances Object Recognition When Perception Is Challenging

We assessed object recognition behavior as a function of difficulty and context using a 2×2 design with factors context (intact vs. scrambled scenes) and difficulty (low vs. high). For this we created forty-eight unique target objects from six high-level categories (Food, Electronics, Containers, Plants, Office Supplies, and Home Decor; Fig. 1A) that were embedded into diverse indoor scenes. To manipulate difficulty, we altered object states: in high-difficulty condition, objects were small, partially occluded, or shown from non-canonical viewpoints. These manipulations often required embedding objects in visually cluttered scenes, meaning that object difficulty and scene complexity were correlated as also occurs naturally. Context was manipulated by embedding each object suitably in either an intact scene, thus featuring appropriate naturalistic spatial and semantic structure (Võ & Wolfe, 2013), or embedding the object in a phase-scrambled version of the same scene, thus retaining low-level image statistics but disrupting meaningful layout (Fig. 1B).

**Figure 1.**
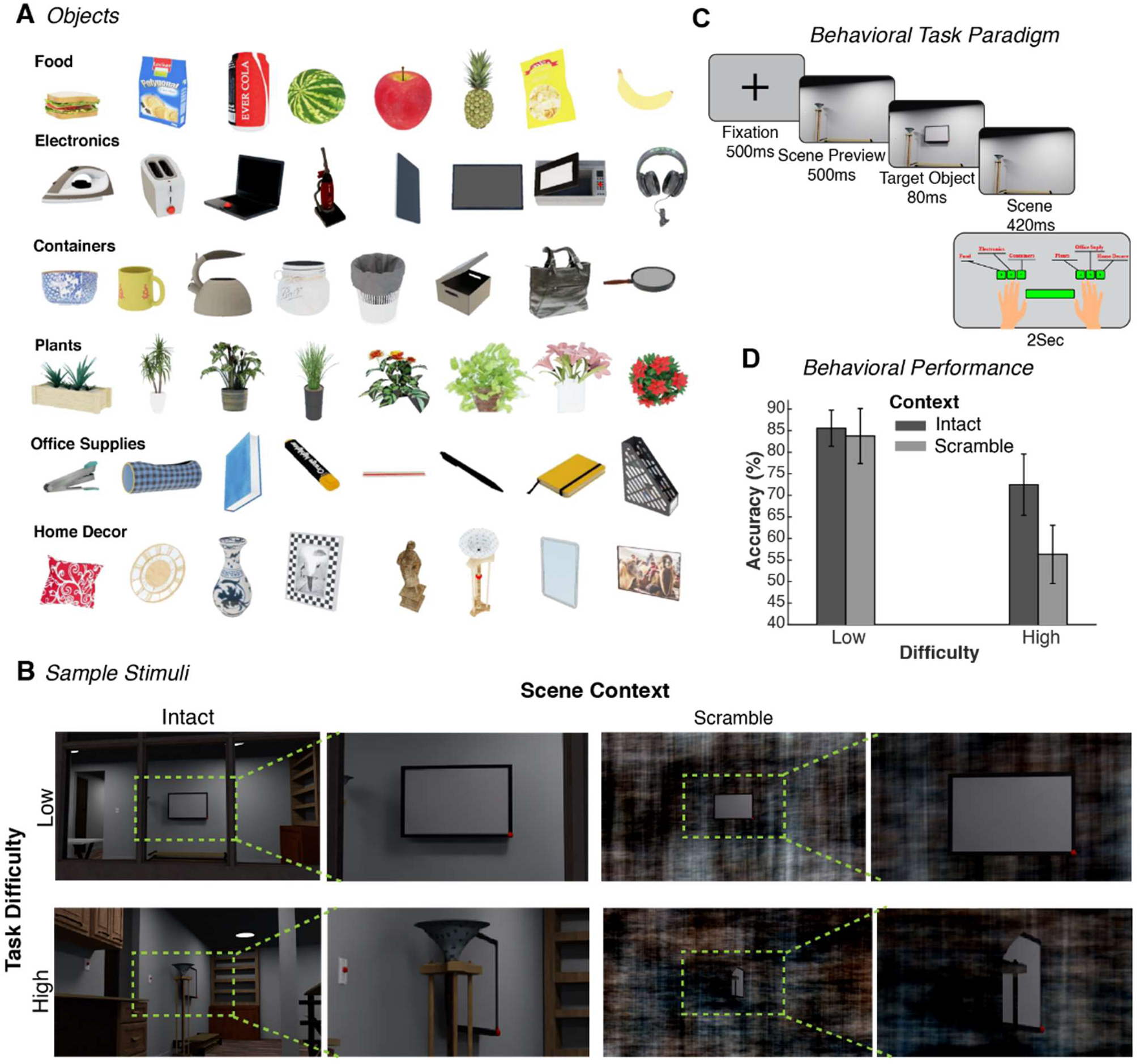
Overview of experimental design and human performance. **A.** Full set of 48 unique objects drawn from six high-level groups: Food, Electronics, Containers, Plants, Office Supplies, and Home Decor. **B.** Sample stimuli illustrating the 2×2 design with two levels of recognition difficulty (low vs. high) and two context types (intact vs. phase-scrambled). Each of the 48 unique objects was embedded into indoor scenes to create both low- and high-difficulty conditions. Contextual structure was manipulated by generating a phase-scrambled version of each background while keeping the target object intact. The final stimuli set included 96 intact images and their 96 corresponding scrambled counterparts. Left images depict original images, right images depict enlarge portions highlighting target objects. **C.** Behavioral task structure. Each trial began with a 500 ms presentation of either an intact scene or scrambled background, followed by an 80 ms presentation of the centrally located target object. Participants categorized the object into one of six high-level groups using a key press (response window: 2 s). **D.** Human recognition accuracy across conditions. The vertical axis reflects categorization accuracy (chance = 16.6%). Bars represent group level mean across 16 participants; error bars indicate standard deviations.

To assess behavior, we used a rapid visual categorization task (N = 16, 20–35 years of age, 6 female). Each trial began with a 500 ms fixation period, followed by a 500 ms preview of the scene (without the target object), followed by an 80 ms presentation of the target object embedded in the scene, and then re-presentation of 420 ms duration of the scene without the object (Fig. 1C). Last, participants categorized the target into one of six high-level groups (e.g., food, electronics, plants) using a button press.

To formally assess the effects of scene context and task difficulty on recognition accuracy, we conducted a non-parametric ANOVA. This analysis revealed significant main effects of both context (F(1, 45) = 70.23, p < 0.001, where F(1, 45) indicates the F-statistic with 1 and 45 degrees of freedom) and difficulty (F(1, 45) = 259.23, p < 0.001), as well as a significant interaction between the two factors (F(1, 45) = 34.55, p < 0.001). To assess the interaction, we used post-hoc tests (sign-rank tests). We found that recognition accuracy was uniformly high when the factor difficulty was low, with no significant difference between intact scenes (85.5%) and scrambled backgrounds (83.7%) (p > 0.05, signed-rank test; Fig. 1D). However, when the factor difficulty was high, overall accuracy declined (p = 0.002), and scene context significantly boosted performance: a significant difference (p = 0.004) between 72% accuracy for intact scenes vs. 56% for scrambled backgrounds.

This shows that meaningful contexts aids object recognition particularly under challenging perceptual conditions, that is when local object features are insufficient to support object recognition on their own due to object degradation or ambiguity.

### Language-Aligned Training Drives Human-Level Accuracy

To determine the extent to which computational models match human context-sensitivity in object recognition, we compared recognition accuracy and context sensitivity between models and human participants. A total of 70 models (Fig. 2A) spanning a wide range of architectures and training paradigms were evaluated using the same stimulus set as in the human experiment.

**Figure 2.**
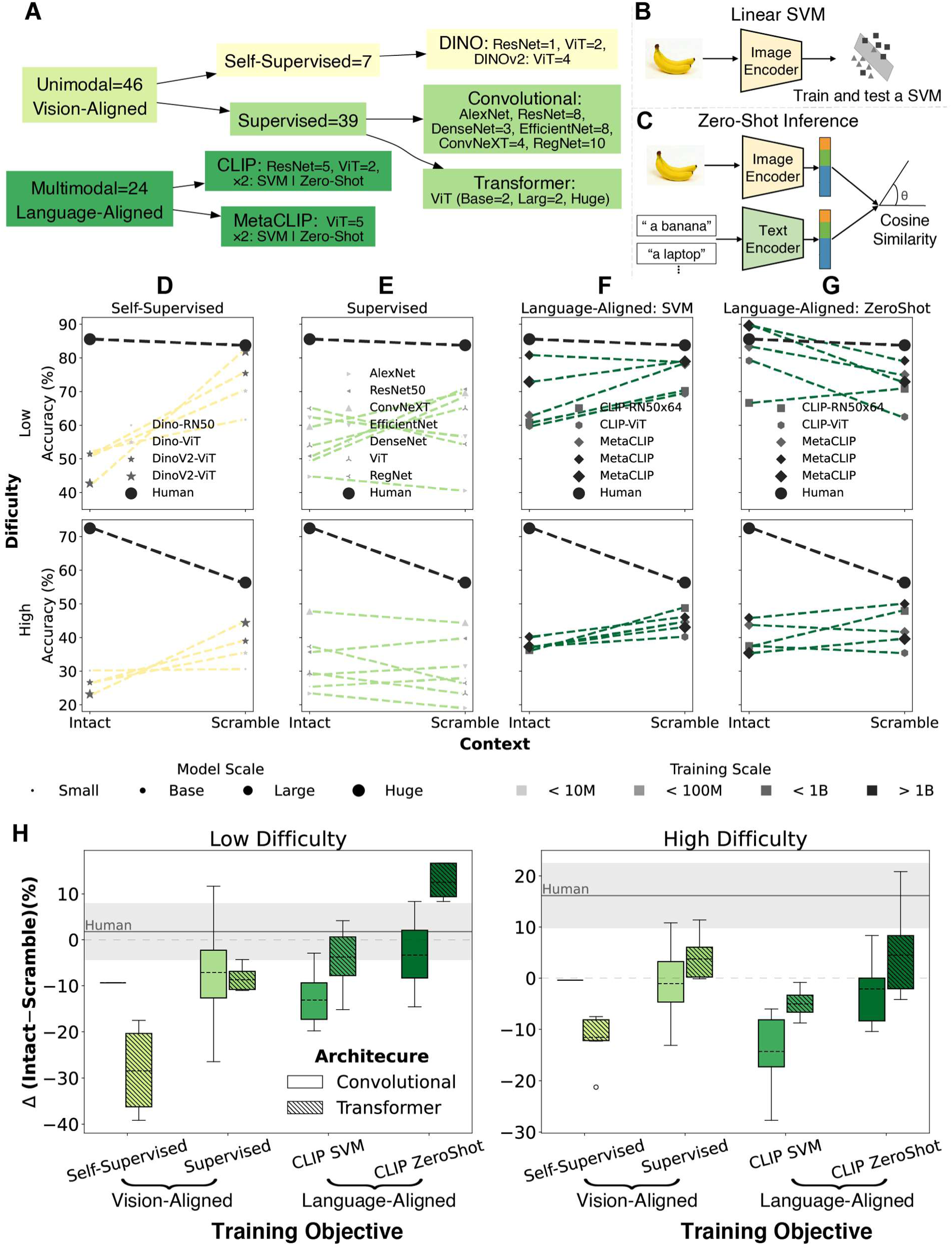
Model accuracies across task conditions and evaluation methods. **A.** Overview of computational models evaluated, grouped by training paradigm (unimodal vs. multimodal), architecture (e.g., convolutional vs. transformer). **B,C.** Evaluation method: (B) Linear-SVM, where a classifier is trained on visual embeddings from the image encoder; (C) Zero-shot, where object category is predicted by computing cosine similarity between visual and text embeddings and selecting the highest match. **D-G.** Object recognition accuracy across context (intact vs. scrambled) and difficulty (low vs. high) for humans and 21 computational models. Subplots are grouped by model type (columns) and difficulty level (rows). Each marker represents a model; marker size indicates parameter count and saturation reflecting training data scale. Human performance is shown as a black circle with a solid line. Dotted lines connect model performance across context; line colors indicate model groups, following the color scheme shown in panel A. **H.** Quantifying sensitivity to contextual structure across models. Boxplots summarize context effects, calculated as the difference in object recognition accuracy between intact scenes and scrambled versions (intact minus scramble), separately for the low- and high-difficulty condition. Models are grouped by training objective, and evaluation method, with architecture type (convolutional vs. transformer) also distinguished within each group. Each boxplot displays the distribution of context effects within a model group. Horizontal lines within boxes indicate group means. Box colors match the color scheme shown in panel A, and distinct hatch patterns are used to differentiate convolutional and transformer architectures. The solid gray line denotes the mean human context effect, with the lighter gray shaded band represents ±1 standard deviation across participants. This visualization reveals how closely different model classes approximate human-like use of context.

We assessed model performance using two complementary evaluation methods that probe distinct aspects of object recognition in models.

First, to assess the discriminative power based on visual representations alone (Fig. 2B), we used linear classification with support vector machines (SVMs). This approach tests whether a model’s visual representations are sufficiently rich to support context-sensitivity. For each model, we trained a SVM on visual embeddings extracted from isolated object images with plain background. Importantly, this method isolates the model’s raw visual representation capacity, independent of language or prior knowledge.

Second, to assess conceptual generalization via language we used zero-shot classification (Fig. 2C). This evaluates the models’ ability to generalize based on language-aligned training, analogous to how humans use semantic knowledge to interpret objects under novel or ambiguous conditions. We evaluated CLIP models without task-specific training by comparing the image embeddings to category-relevant text prompts (e.g., “a photo of a banana”) using cosine similarity.

In both cases, model predictions from the 48 object categories were mapped onto the six behavioral categories used in the human task, enabling direct comparison with human performance across conditions (as summarized in Fig. 1D).

Fig. 2D-G presents the results for a subset of 21 representative models, selected to reflect general patterns across the diversity of training paradigms, architectures, data scale, and evaluation methods investigated in full (for complete results see Supplementary Table 1; for model group summaries see Supplementary Fig. 1). These models include unimodal supervised models (Fig. 2E, e.g., CNNs trained on ImageNet), unimodal self-supervised models (Fig. 2D, e.g., trained with contrastive learning), and multimodal language-aligned models (e.g., CLIP trained on image-text pairs), evaluated via either linear classification (Fig. 2F) or, zero-shot inference (Fig. 2G).

We compared model and human performance in accuracy across the key experimental factors of context and difficulty.

Classic unimodal vision models (Fig. 2E) trained in supervised fashion—including CNNs and Vision Transformers (ViT)—performed significantly worse than humans across all conditions (11–50% deficits; ps < 0.001,, indicating all p-values were below 0.001, rank-sum test; see Supplementary Fig. 3 for full p-value results). We speculate that these models failed to leverage scene context when recognizing ambiguous or occluded objects, likely due to their training on ImageNet (up to 14M labeled images), which lacks the semantic structured and contextual diversity typical of real-world environments (Geirhos et al., 2020; Ullman et al., 2016; Zhou et al., 2019).

Unimodal self-supervised learning (SSL) models (Fig. 2D) such as DINO and DINOv2 (Oquab et al., 2023), trained on much larger datasets (e.g., 142M images), still fell short of human performance. In intact scenes, they underperformed humans by 27–49% (ps < 0.001). Notably, performance improved relatively in the scrambled condition: for example, DINOv2 nearly matched human accuracy (81.9% vs. 83.8%, p > 0.05) in the low-difficulty scrambled condition. This pattern suggests that large SSL models treat structured scene background here as noise rather than as sources of useful contextual information, and thus do not benefit from meaningful scene structure.

Multimodal language-aligned models, trained on large-scale image-text data using both linear (Fig. 2F) and zero-shot classification (Fig. 2G) achieved performance comparable to humans. Notably, ViT-based VLMs in zero-shot setting performed similar to human (p > 0.05; see Supplementary Fig. 1 and Supplementary Table 2). This suggests that language-aligned training enables near-optimal category-level recognition when visual conditions are of low difficulty. Importantly, this human-level performance appears to reflect not only the benefits of semantic supervision, but also the architectural strengths of ViT backbones. As elaborated in Supplementary Discussion 1, vision transformers—particularly when paired with language-aligned training—offer a distinct advantage for context-rich recognition, highlighting a synergistic interaction between architecture and supervision type.

However, in the high-difficulty condition, where objects were smaller, occluded, or presented from atypical viewpoints, MetaCLIP and other CLIP models still underperformed relative to humans. This indicates that these models align in performance only under low perceptual difficulty, while they fail to fully match human robustness when object recognition is demanding. One contributing factor may be that, in the human experiment, participants previewed the scene before the object appeared, a temporal structure that likely facilitated attentional guidance and scene interpretation in difficult trails, that is not captured by current model inference procedures.

In sum, the model comparison revealed that unimodal vision models fall short in matching human object recognition accuracy across difficulty levels, whereas multimodal language-aligned models rivaled human object recognition performance when perceptual difficulty was low.

### VLMs When Probed Using Zero Shot Inference show similar performance in the low difficulty condition

Humans flexibly leverage scene context to support recognition under visual uncertainty (Fig. 1D). Participants showed strong positive context benefit in the high-difficulty condition, with a +16.4% gain in intact scenes compared to scrambled versions (*p* < 0.001). In contrast, in the low-difficulty condition, where recognition was relatively easy, context effects were minimal (+1.8%, *p* > 0.05). This pattern reflects adaptive behavior: contextual cues are recruited when local features are insufficient and otherwise ignored.

We quantified model context sensitivity using the same metric as in humans – i.e., the accuracy difference between intact and scrambled conditions (Fig. 2H) for all 70 models, and show the results grouped by model kind and inference procedure.

In the low difficulty condition, all models with the exception of VLMs with transformer-based architecture evaluated using zero-shot classification showed a negative or non-significant context effect, rather than a human-like positive context effect (SSL models: −28.4% for ViT, −9.2% for ResNet; supervised models: −8.4% for ViT, −7.1% for CNNs; and VLMs evaluated using linear classification: −12.8% for ViT, −3.9% for ResNet; see Supplementary Table 3 for detailed statistical results). This suggests that coherent scenes impaired rather than aided object recognition. In contrast, VLMs with ViT architecture evaluated using zero-shot classification that showed a human-like positive context effect of (+12.7%, p = 0.016), while their ResNet-based counterparts exhibited no significant context effect (−3.2%, p > 0.05) with the difference between architectures reaching significance (p = 0.006).

In the high-difficulty condition, SSL models and VLMs when probed using linear classification showed negative context effects (SSL: −11.4% for ViT, −0.35% for ResNet; VLMs: −14.2% for ResNet and −4.9 for ViT; all *ps* < 0.01). Supervised models and VLMs probed with zero-shot classification had no significant context effects (ViT-based models: +4.3% and +4.7% respectively; and ResNet-based models: −1%, −2%; ps > 0.05), showing an improved but still incomplete fit to human behavior. Also, all models had lower context sensitivity than humans (all ps< 0.05). Together this highlights ViT-based VLMs probed with zero-shot inferences as the modelling procedure incompletely, yet most closely matching human context-sensitive behavior.

## Discussion

Humans excel at recognizing objects even in cluttered, ambiguous environments—an ability that artificial vision systems still struggle to match. This capacity is thought to reflect the brain’s ability to integrate incoming sensory information with prior knowledge, including semantic expectation and contextual cues (Lupyan et al., 2020; Peelen et al., 2024). Here, we investigated whether VLMs, trained on large-scale image-text datasets, can approximate this kind of context-sensitive object recognition. Our findings yield two key insights.

First, multimodal training—particularly with natural language—enables models to better approximate human-level performance than unimodal training. One likely reason is that language supervision fosters semantically structured visual representations, which support more flexible and context-sensitive inference. This interpretation is consistent with prior work (Conwell et al., 2024; Doerig et al., 2024) and is further supported by our supplementary analysis (Supplementary Discussion 1), which shows that recognition accuracy in coherent scenes increases systematically with the semantic richness of supervision—with VLMs outperform models trained on larger but non-aligned datasets. This effect is particularly pronounced for ViT-based VLMs, which consistently outperform convolutional architectures. These results highlight the importance of language alignment—not just training scale—for achieving human-like context sensitivity in object recognition.

Importantly, the benefits of language supervision may not arise solely from explicit language exposure. Although VLMs are trained on image–text pairs, emerging evidence suggests that representational convergence between language and vision may reflect shared sensitivity to environmental structure. For instance, Conwell et al. (2025) found that both VLMs and language models predict visual cortical activity in non-human primates—despite the latter’s lack of language exposure. This suggests that the representational overlap may stem from internalizing the statistical structure of the visual world, rather than from language per se.

While this perspective highlights representational convergence, our results points to a second, complementary insight: inference mechanisms matter. Language plays a critical role not only in shaping representations but also in guiding flexible interpretation. While VLMs trained with language supervision encode rich, object discriminative representations, these alone do not yield human-like context sensitivity. The similarity to human behavior improves when inference explicitly incorporate language—such as in zero-shot setups where image embeddings are directly matched to text prompts, rather than decoded via fixed classifier.

Zero-shot inference allows models to adapt their interpretation of visual input based on natural language cues that reflect both object identity and scene-specific expectations. This offers a computational mechanism for context-sensitive recognition—mirroring human vision’s integration of bottom-up sensory input with top-down conceptual knowledge (Lupyan et al., 2020; Peelen et al., 2024).

This insight contributes to a broader debate on comparing artificial and biological systems (Cichy et al., 2019; Schrimpf et al., 2018). While many studies emphasize aligning models with the brain in terms of architecture, training data, or representational similarity; our findings show that inference procedures must also be considered. CLIP’s ability to approximate human-like context sensitivity depends on both multimodal training and language-guided inference. Without the latter, key behavioral signatures are lost. Thus, meaningful comparisons between artificial systems and human cognition must align not only their learned representations, but also the interpretive procedures through which they are applied.

We note, however, that for all models and evaluation metrics the fit to human context sensitivity was incomplete. In particular in the high difficulty condition no model matched human behavior. One likely contributing factor is the temporal structure of the behavioral experiment: participants previewed the entire scene briefly before the target object appeared—which may have facilitated scene-based expectations—especially under challenging conditions. Another limitation lies in the constrained input resolution of most models (typically 224×224 pixels), which may hinder their ability to detect small or partially visible objects in cluttered scenes.

To bridge these gaps, future work could explore emerging vision-language models capable of processing images sequences or modeling temporal dynamics, thereby better approximating the temporal dynamics of human visual perception. Additionally, leveraging models that support higher-resolution inputs or process wider fields of view may enhance performance in complex, context-rich scenes.

### Modeling Contextual Recognition: Methodological Advances

Beyond the immediate results our work offers methodological advances by providing a framework to compare humans and models with respect to context sensitivity. We thus provide the stimulus material and human behavioral data openly for future research.

To meaningfully compare human and model performance in object recognition within complex scenes, our study aimed to balance ecological validity with experimental control.

We sought to evaluate context-sensitive object recognition using stimuli and conditions that reflect the richness of real-world visual experience. Specifically, we used 3D-rendered indoor scenes from the Omnigibson platform, which offer realistic lighting, depth, occlusion, and object arrangement—features that surpass traditional static or hand-crafted image set in capturing everyday visual complexity.

To isolate the role of semantic scene structure, we compared recognition performance in intact scenes to that in phase-scrambled versions of the same images. This manipulation preserves low-level visual statistics while disrupting global structure and meaning, enabling us to assess how coherent context facilitates recognition beyond simple background removal (e.g., blank or uniform fields).

We also systematically varied task difficulty using naturalistic factors such as occlusion, object size, lighting, and clutter—common challenges in daily visual perception. These manipulations allowed us to probe recognition under both easy and difficult conditions while maintaining ecological relevance.

Crucially, our design incorporated a brief temporal separation between the presentation of the scene and the appearance of the target object. This mimics real-world viewing conditions, where observer typically perceives the scene layout before focusing on specific items. Beyond ecological validity, this temporal structure may engage top-down expectations, facilitating the integration of incoming sensory input with prior semantic knowledge. This approach builds on prior findings that scene previews can enhance subsequent object recognition (Chen et al., 2022; Võ & Wolfe, 2013), and offers a useful framework for studying the dynamics of context integration under temporally constrained viewing.

Together, these methodological choices allowed us to examine contextual facilitation under challenging yet structured conditions, supporting direct comparisons between human behavior and model predictions in the context of naturalistic visual inference.

## Methods

### Scene-Object Dataset

To investigate how scene context influences object recognition, we developed a dataset that systematically manipulates object-scene relationships while preserving naturalistic structure. Real-world images exhibit rich semantic and spatial regularities, but their complexity makes it difficult to isolate factors such as contextual congruency or visual clutter. To address this, we used the OmniGibson platform—an embodied AI environment offering fine-grained control over naturalistic visual stimuli (Ge et al., 2024; Li et al., 2023).

OmniGibson contains over 50 scene types (e.g., homes, offices, hotels, schools, restaurants) and a large library of 3D objects that can be placed in semantically appropriate locations. We selected 48 object categories spanning six high-level semantic classes: food, electronics, containers, plants, office supplies, and home decor. These were based on metadata from the THINGSplus dataset (Hebart et al., 2023), which groups 1,825 everyday objects into 53 semantic categories. From each of our six categories, we selected eight representative objects (e.g., laptop, vacuum, iron for electronics) and placed them in semantically and spatially congruent locations using OmniGibson’s scripting tools to ensure ecological validity.

To introduce controlled variation, we manipulated camera distance and angle, lighting conditions, object visibility, and overall scene complexity. This yielded images at two levels of difficulty: low (e.g., minimal clutter, higher visibility) and high (e.g., increased occlusion and crowding). Each of the 48 unique objects appeared in both conditions, producing 96 total images (48 per difficulty level; Fig. 1A–B). All images were converted to grayscale to remove color salience and isolate structural and semantic cues.

#### Scrambled Backgrounds

To disrupt contextual structure while preserving low-level image properties, we generated phase-scrambled versions of each scene. This manipulation degrades perceptual coherence while retaining basic image statistics, allowing us to isolate the influence of global structure on recognition.

Phase scrambling was performed in the Fourier domain. We extracted each image’s amplitude and phase spectra using a Fast Fourier Transform (FFT), replaced the phase spectrum with a randomized version (scaled by a scrambling factor, p ∈ [0,1]), and reconstructed the image using the inverse FFT. This procedure preserved global contrast and spatial frequency contrast while destroying semantic and spatial structure. The target object remained intact, but the surrounding context became incoherent. Scrambled images were generated using validated code from Hebart (2009) (http://martin-hebart.de/webpages/code/stimuli.html).

### Behavioral Experiment

To assess how humans use context to recognize objects, we conducted a behavioral experiment using the scene-object images. The goal was to test whether structured scene context facilitates object recognition under perceptual constraints.

Sixteen participants (7 female, 9 male; age 20–35), all with normal or corrected-to-normal vision, were recruited from the university subject pool.

We used a sequential scene-object presentation paradigm (Chen et al., 2022) to temporally separate the scene and target minimizing masking and isolating semantic facilitation.

Each trial began with a 500 ms fixation cross, followed by a 500 ms preview of the scene without the target object. The target object then appeared briefly for 80 ms, centrally embedded in the same scene, followed by 420 ms of the unchanged scene. Participants categorized the target object using one of six predefined keys corresponding to the six high-level groups (see Fig. 1C).

Trial order was randomized, and all conditions (intact vs. scrambled; low vs. high difficulty) were fully counterbalanced. Before the main task, participants completed a training phase using plain-background objects to learn category-to-key mappings.

The full experiment included 192 trials (48 objects × 2 contexts × 2 difficulty levels). Images subtended 21° × 16° of visual angle, with objects no larger than 4°. The experiment was implemented in MATLAB using Psychophysics Toolbox (Brainard, 1997).

### Computational Models

To evaluate whether artificial models exhibit human-like context sensitivity, we tested 70 models across four families: unimodal vision-aligned DNNs like CNNs, Vision Transformers (ViTs), and SSL models, and multimodal language-aligned models such as CLIP (Fig. 2A). All models were evaluated on the same six high-level categories used in the behavioral task and tested under the same four experimental conditions (intact/scramble x low/high) To approximate foveal input, images were cropped to the central 6° of visual angle and resized to 224×224 pixels, and converted to grayscale.

#### Training and Evaluation

For models requiring supervised training (e.g., SVMs), we used a separate plain-background object dataset from the human training phase. We training classifiers using 10 random 80/20 train-validation splits. For CLIP models, we used two methods:

- **SVM classification** using extracted visual embeddings.
- **Zero-shot classification**, where image embeddings were compared to category text prompts (e.g., “a photo of a laptop”) using cosine similarity.

This unified framework ensured that all models were tested under conditions directly comparable to the human task.

#### Model Families CNNs

We evaluated standard convolutional networks including AlexNet, ResNet (18–152 layers), DenseNet, RegNet, EfficientNet, and ConvNeXT. All models were pretrained on ImageNet-1k or 21k. These architectures build hierarchical features through convolutional layers and residual connections (He et al., 2015; Krizhevsky et al., 2012).

#### ViTs

ViTs use self-attention to model global relationships between image patches using self-attention mechanisms without convolution (Dosovitskiy et al., 2020). We used ViTs pretrained on ImageNet and classified using SVMs on extracted features.

#### SSL Models

We tested DINO and DINOv2, which learn visual representations via self-distillation and contrastive learning (Caron et al., 2021; Oquab et al., 2023). DINO was trained on ImageNet-1k; DINOv2 used 142M unlabeled images.

#### VLMs

We evaluated CLIP (Radford et al., 2021), which jointly learns visual and textual embeddings from large-scale image–text pairs. We used both OpenAI CLIP (trained on 400M pairs) and MetaCLIP (400M and 2.5B pairs; Xu et al. (2023)). Each was tested via both SVM and zero-shot classification.

#### Human-Model Comparison

In the behavioral task, humans categorized objects into one of six high-level categories via keypress. Models, by contrast, initially classified images into one of 48 fine-grained object categories. We then mapped model predictions to their corresponding high-level categories to align the format with human responses. This reflects the idea that humans likely perform fine-grained recognition prior to categorical abstraction. While the response formats differed, this mapping enabled a controlled comparison evaluation of context sensitivity across human and models.

### Statistical Testing

For behavioral data, we conducted a 2×2 repeated-measures non-parametric ANOVA in R to examine the effects of context (intact vs. scramble) and difficulty (low vs. high) on recognition accuracy. Post-hoc comparisons were conducted using Wilcoxon signed-rank tests (Wilcoxon, 1945) for pairwise contrast.

For model data, context effects were computed as the accuracy difference between intact and scrambled conditions. We compared these context effect scores across model families using Wilcoxon rank-sum tests.

For SVM-based models, we report accuracy distributions across 10 random training splits. In contrast, zero-shot CLIP models yield a single accuracy score per condition; we acknowledge that this limits variance estimation and restricts robust statistical comparisons for these models.

## Supplementary

### Supplementary Discussion 1: What drives variation in model performance across natural scenes?

**Supplementary Fig. 1.**
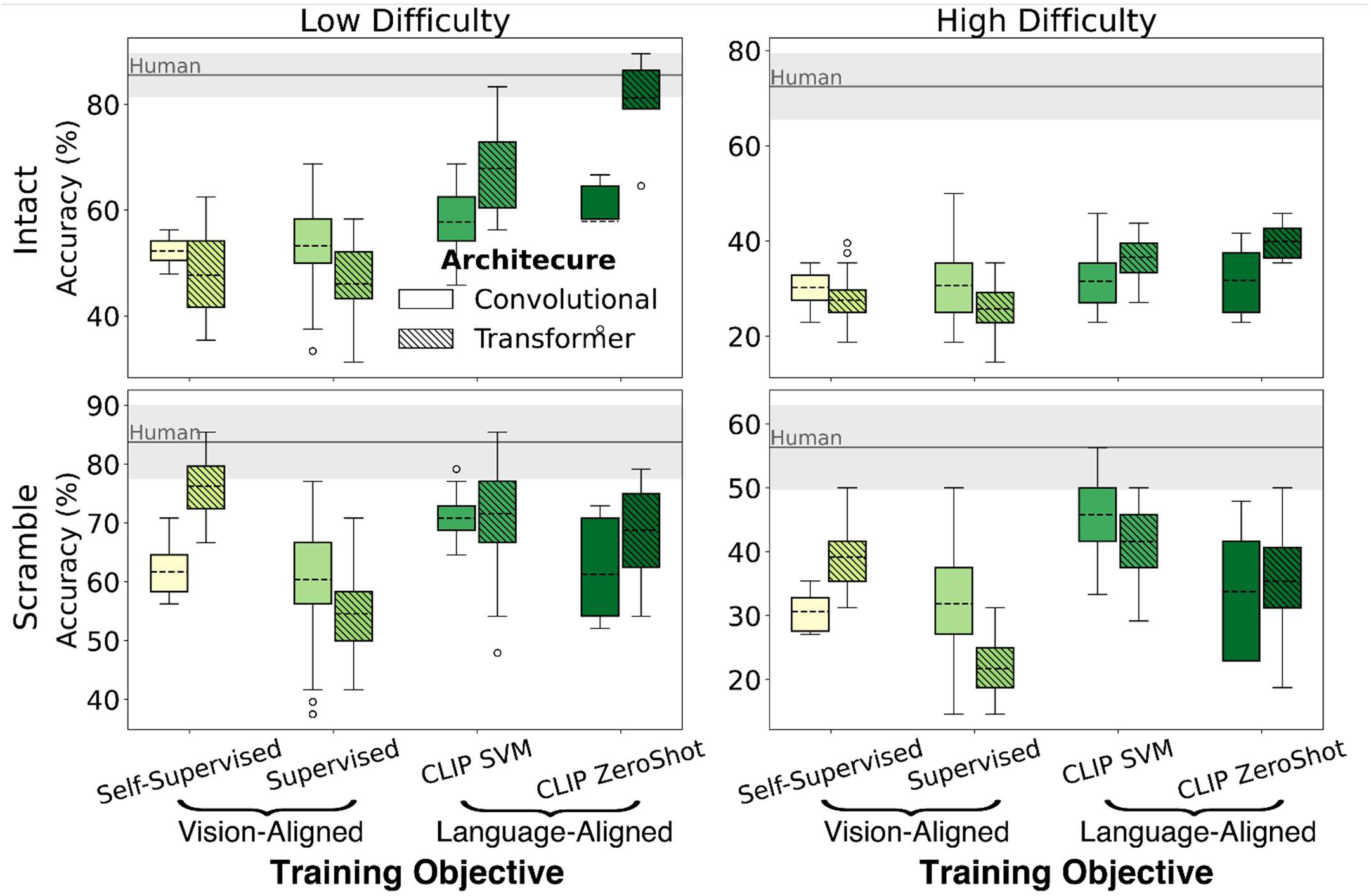
Relationship between training objective and performance across scene context and difficulty. The 2×2 subplot layout displays performance across levels of scene context (intact vs. scrambled; rows) and task difficulty (low vs. high; columns). Within each subplot, models are grouped by training objective (e.g., supervised, self-supervised, language-supervised), with boxplot hatch patterns indicating architecture type (convolutional vs. transformer) and box colors corresponding to model groups shown in Figure 2. Dashed horizontal lines indicate group means within each subplot. The solid gray line denotes the mean human accuracy, with the lighter gray shaded band represents ±1 standard deviation across participants.

What factors best explain the variation in model performance across naturalistic scenes? We examined three key factors: (1) the nature of training supervision, (2) the architecture of the vision backbone, and (3) the scale and diversity of training data—each of which has been previously implicated in driving model performance and brain-like representations (Wang et al., 2024; Conwell et al., 2025; Doerig et al., 2024; Radford et al., 2021).

Models were first grouped on an ordinal scale by supervision type: self-supervised (which learn without category labels, e.g., DINO), category-supervised (trained on labeled object categories, e.g., ResNet and ViT), and multimodal language-aligned (e.g., CLIP).

Recognition accuracy increased systematically with the semantic richness of supervision. As shown in Supplementary Fig. 1, models trained with stronger semantic supervision— especially VLMs—achieved the highest accuracy in intact natural scenes. A linear regression across supervision types revealed a strong correlation between semantic supervision and accuracy for intact natural scenes (Pearson’s r = 0.99, p = 0.008, two-tailed t-test), but not for scramble scenes.

**Supplementary Fig. 2.**
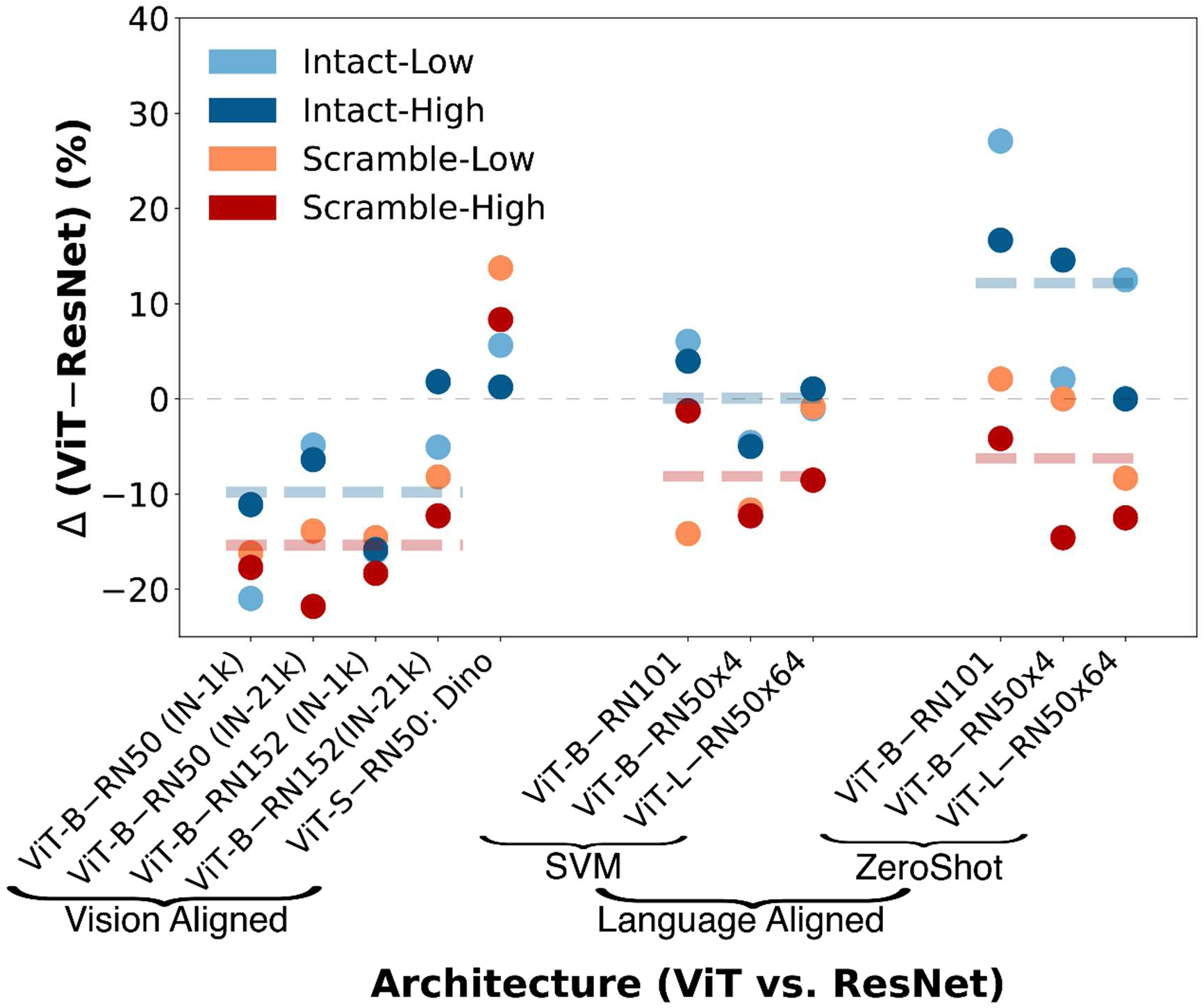
**Performance Comparison Across Architecture.** Accuracy difference between Vision Transformers (ViT) and ResNet (CNN) architectures, matched for training dataset (e.g., ImageNet or 400M image-text pairs), parameter counts, training paradigm, and evaluation methods (e.g., SVM or Zero-Shot). Each circle represents the accuracy difference between a ViT and a ResNet model. Dashed lines indicate the mean accuracy differences across architecture pairs within each model group for the intact (blue) and scramble (red) conditions.

Next, we assessed the contribution of vision backbone architecture. As shown in Supplementary Fig. 1, ViT-based VLMs outperformed ResNet-based VLMs in the intact, low-difficulty condition (81.8% vs. 58.4% for zero-shot, p = 0.01; 67.2% vs. 57.9% SVM, p = 0.048; Supplementary Table 2), suggesting the architectural advantage of ViT. To isolate architectural effects, we curated a subset of models matched on training data (e.g., ImageNet or 400M image-text pairs), parameter counts, and training paradigms (vision-aligned vs. language-aligned; see Supplementary Fig. 2). This allowed us to directly assess architectural differences between ViT and ResNet backbones, independent of confounding factors.

Among vision-aligned models and VLMs evaluated using SVM classification, ResNet architectures generally outperformed ViTs. In vision-aligned group, ResNets yielded higher accuracy in both intact (−9.8%, p = 0.015) and scrambled scenes (−15.2%, p = 0.007), with the exception of DINO—a self-supervised ViT that exceeded ResNet performance (+3.43% and +11.04 for intact and scramble respectively). Similarly, in language-aligned group, ResNets outperformed ViTs in scrambled scenes (−7.9%, p = 0.03), and performed similarly in intact scenes (0.2%, p > 0.05). There was also a marginal interaction between architecture and scene structure (p = 0.06).

In contrast, when language-aligned models were evaluated using zero-shot inference, ViTs significantly outperformed ResNets in intact scenes (+12.4%, p = 0.04). This pattern reversed in scrambled scenes, where ResNets showed a modest advantage (−6.1%; p = 0.08). The interaction between architecture and scene structure was significant (p = 0.03), indicating a scene-dependent advantage for ViTs.

These findings suggest that ViT offers a distinct advantage for zero-shot inference in coherent scenes—likely due to their global self-attention mechanisms, which better capture structured scene context than the local feature emphasis of CNNs (Raghu et al., 2021; Nasser et al., 2021). This advantage may also reflect stronger visual-language alignment, likely stemming from their shared transformer-based architecture across vision and language components. Consistent with this interpretation, results from CLIP (Radford et al. 2021) and MetaCLIP (Xu et al., 2023) show that ViT-based VLMs outperformed convolutional backbones, further underscoring the central role of transformers in high-performing VLMs.

Finally, we evaluated the impact of data scale and diversity, ordering models by the number of training examples: category-supervised models were trained on ≤14 million labeled images, self-supervised models were trained on up to 142 million unlabeled images, and language-aligned models on as many as 2.5 billion multimodal image–text pairs. While larger datasets were modestly associated with higher accuracy (Pearson r = 0.27, p < 0.001), data scale alone did not account for performance differences. Notably, self-supervised models trained on massive datasets still underperformed compared to VLMs, highlighting that semantic alignment, not just data scale, is essential for robust recognition.

In summary, these findings highlight that semantic supervision is the primary driver of model performance in context-sensitive recognition, followed by architectural design and training data diversity. Importantly, ViT backbones—particularly when paired with language-aligned training—are especially well suited for capturing structured visual context and supporting high-performance zero-shot recognition.

**Supplementary Table 1.** Object recognition accuracy (± standard deviation) for all evaluated models across four experimental conditions. Each row reports results for one model, showing accuracy under two context conditions (Intact, Scrambled) and two difficulty levels (Low, High). Models are grouped by evaluation type (SVM or Zero-Shot). For SVM-based evaluations, the reported values reflect the mean and standard deviation over 10 random train/test splits. For Zero-Shot evaluations, only a single accuracy value is reported per condition (standard deviations are not applicable).

| Model Name | Low-Intact | Low-Scramble | High-Intact | High-Scramble | Eval Type | Training Scale (Millions) | Parameters (Millions) |
| --- | --- | --- | --- | --- | --- | --- | --- |
| Dino-ResNet50 | 52.2 ± 2.3 | 61.6 ± 4.3 | 30.2 ± 3.7 | 30.6 ± 2.8 | SVM | 1 | 23 |
| Dino-ViT<sub>s</sub>8 | 57.9 ± 2.2 | 75.4 ± 3.2 | 31.4 ± 4.9 | 38.9 ± 2.6 | SVM | 1 | 21 |
| Dino-ViT<sub>b</sub>8 | 51.0 ± 3.1 | 70.2 ± 2.2 | 26.7 ± 3.8 | 35.4 ± 2.6 | SVM | 1 | 85 |
| DinoV2-ViT<sub>s</sub>14 | 39.4 ± 2.7 | 73.8 ± 3.0 | 31.0 ± 3.3 | 39.0 ± 3.0 | SVM | 142 | 21 |
| DinoV2-ViT<sub>b</sub>14 | 44.0 ± 2.0 | 80.8 ± 2.0 | 26.7 ± 2.4 | 38.5 ± 4.2 | SVM | 142 | 86 |
| DinoV2-ViT<sub>l</sub>14 | 51.5 ± 1.9 | 75.4 ± 4.0 | 26.7 ± 3.3 | 39.0 ± 3.4 | SVM | 142 | 300 |
| DinoV2-ViT<sub>g</sub>14 | 42.7 ± 1.9 | 81.9 ± 2.3 | 23.1 ± 2.4 | 44.4 ± 3.7 | SVM | 142 | 1100 |
| AlexNet | 44.7 ± 3.2 | 40.6 ± 2.7 | 23.3 ± 1.7 | 19.0 ± 2.1 | SVM | 1 | 60 |
| ResNet18-v1 | 49.4 ± 3.2 | 62.1 ± 2.0 | 30.8 ± 2.6 | 39.0 ± 3.7 | SVM | 1 | 11 |
| ResNet34-v1 | 52.9 ± 3.1 | 57.5 ± 2.7 | 24.4 ± 2.1 | 37.3 ± 4.3 | SVM | 1 | 21 |
| ResNet50-v1 | 56.0 ± 2.9 | 61.8 ± 2.7 | 32.4 ± 3.4 | 38.8 ± 3.6 | SVM | 1 | 25 |
| ResNet101-v1 | 57.9 ± 2.2 | 58.8 ± 5.1 | 37.7 ± 3.5 | 27.9 ± 4.1 | SVM | 1 | 45 |
| ResNet152-v1 | 51.0 ± 2.1 | 60.2 ± 2.2 | 37.1 ± 2.8 | 39.4 ± 3.8 | SVM | 1 | 60 |
| ResNet50-v2 | 50.8 ± 2.4 | 70.7 ± 3.1 | 35.7 ± 2.8 | 39.7 ± 2.8 | SVM | 14 | 25 |
| ResNet101-v2 | 52.9 ± 2.1 | 63.5 ± 2.3 | 31.3 ± 2.5 | 30.6 ± 2.8 | SVM | 14 | 45 |
| ResNet152-v2 | 51.0 ± 4.5 | 65.0 ± 6.8 | 27.5 ± 2.9 | 30.2 ± 1.7 | SVM | 14 | 60 |
| ConvNeXT-T | 42.7 ± 3.0 | 65.0 ± 4.6 | 31.3 ± 2.6 | 36.9 ± 3.0 | SVM | 1 | 27 |
| ConvNeXT-S | 52.1 ± 3.7 | 64.0 ± 2.3 | 36.0 ± 3.1 | 37.1 ± 3.5 | SVM | 1 | 50 |
| ConvNeXT-B | 60.0 ± 3.3 | 67.5 ± 1.9 | 28.3 ± 4.0 | 39.2 ± 4.6 | SVM | 1 | 86 |
| ConvNeXT-L | 59.6 ± 2.1 | 69.6 ± 3.4 | 47.7 ± 2.5 | 44.4 ± 3.1 | SVM | 1 | 230 |
| EfficientNet-B0 | 59.6 ± 3.1 | 62.5 ± 2.6 | 33.1 ± 3.0 | 23.8 ± 2.5 | SVM | 1 | 5 |
| EfficientNet-B1 | 53.1 ± 3.4 | 51.7 ± 3.6 | 33.8 ± 2.6 | 26.0 ± 4.0 | SVM | 1 | 8 |
| EfficientNet-B2 | 60.0 ± 3.1 | 62.1 ± 5.5 | 30.0 ± 5.1 | 21.5 ± 2.1 | SVM | 1 | 9 |
| EfficientNet-B3 | 51.3 ± 3.1 | 58.5 ± 2.7 | 23.5 ± 4.4 | 24.2 ± 1.7 | SVM | 1 | 12 |
| EfficientNet-B4 | 55.6 ± 3.7 | 61.9 ± 3.0 | 24.0 ± 3.5 | 28.8 ± 3.6 | SVM | 1 | 19 |
| EfficientNet-B5 | 56.0 ± 3.0 | 64.0 ± 3.0 | 32.5 ± 2.8 | 24.2 ± 1.7 | SVM | 1 | 30 |
| EfficientNet-B6 | 53.5 ± 3.9 | 51.7 ± 3.7 | 32.1 ± 4.2 | 36.5 ± 3.9 | SVM | 1 | 43 |
| EfficientNet-B7 | 62.3 ± 3.3 | 56.7 ± 6.2 | 29.0 ± 2.5 | 31.5 ± 3.4 | SVM | 1 | 66 |
| DenseNet121 | 43.3 ± 3.8 | 69.8 ± 3.9 | 22.5 ± 2.4 | 35.6 ± 0.6 | SVM | 1 | 7 |
| DenseNet169 | 49.6 ± 5.6 | 68.1 ± 3.7 | 25.4 ± 3.9 | 27.9 ± 1.9 | SVM | 1 | 13 |
| DenseNet201 | 41.7 ± 6.7 | 61.5 ± 5.1 | 20.8 ± 1.6 | 24.8 ± 1.5 | SVM | 1 | 20 |
| ViT-B-v1 | 35.0 ± 2.8 | 45.6 ± 2.9 | 21.3 ± 4.2 | 21.0 ± 3.3 | SVM | 1 | 86 |
| ViT-L-v1 | 47.1 ± 3.5 | 54.0 ± 3.0 | 28.1 ± 3.6 | 26.9 ± 2.4 | SVM | 1 | 304 |
| ViT-B-v2 | 46.0 ± 3.4 | 56.8 ± 3.0 | 29.3 ± 3.7 | 17.9 ± 3.2 | SVM | 14 | 86 |
| ViT-L-v2 | 54.0 ± 2.4 | 65.0 ± 4.6 | 29.4 ± 1.1 | 23.3 ± 2.0 | SVM | 14 | 304 |
| ViT-H-v2 | 47.5 ± 4.0 | 51.8 ± 3.2 | 21.3 ± 2.4 | 21.4 ± 2.5 | SVM | 14 | 632 |
| RegNet-y-400mf-v1 | 48.3 ± 2.2 | 63.1 ± 1.9 | 25.8 ± 3.1 | 28.5 ± 2.3 | SVM | 1 | 5 |
| RegNet-y-400mf-v2 | 44.8 ± 1.7 | 57.5 ± 2.8 | 29.4 ± 3.7 | 32.9 ± 4.3 | SVM | 14 | 5 |
| RegNet-y-1-6gf-v1 | 53.8 ± 3.3 | 59.4 ± 5.3 | 32.7 ± 5.3 | 33.5 ± 3.2 | SVM | 1 | 11 |
| RegNet-y-1-6gf-v2 | 56.5 ± 2.9 | 69.0 ± 3.0 | 36.9 ± 3.5 | 34.0 ± 3.9 | SVM | 14 | 11 |
| RegNet-y-8gf-v1 | 57.5 ± 3.3 | 62.7 ± 2.2 | 28.1 ± 4.1 | 35.2 ± 5.1 | SVM | 1 | 39 |
| RegNet-y-8gf-v2 | 58.5 ± 2.5 | 61.5 ± 2.7 | 37.9 ± 3.7 | 32.1 ± 4.2 | SVM | 14 | 39 |
| RegNet-y-16gf-v1 | 55.6 ± 4.1 | 60.4 ± 1.6 | 25.8 ± 2.7 | 32.7 ± 3.1 | SVM | 1 | 83 |
| RegNet-y-16gf-v2 | 53.1 ± 4.4 | 41.5 ± 3.4 | 28.5 ± 1.9 | 29.4 ± 2.0 | SVM | 14 | 83 |
| RegNet-y-32gf-v1 | 54.2 ± 3.6 | 53.8 ± 1.8 | 32.5 ± 2.7 | 33.5 ± 2.9 | SVM | 1 | 145 |
| RegNet-y-32gf-v2 | 65.0 ± 2.0 | 54.4 ± 5.9 | 37.3 ± 4.5 | 26.5 ± 3.5 | SVM | 14 | 145 |
| CLIP-RN101 | 54.4 ± 4.0 | 70.4 ± 1.8 | 29.2 ± 3.2 | 35.2 ± 1.1 | SVM | 400 | 56 |
| CLIP-RN50 | 52.1 ± 3.6 | 69.4 ± 2.6 | 25.4 ± 2.2 | 53.1 ± 1.9 | SVM | 400 | 38 |
| CLIP-RN50x4 | 65.0 ± 2.2 | 67.9 ± 2.7 | 38.1 ± 3.0 | 46.3 ± 4.7 | SVM | 400 | 86 |
| CLIP-RN50x16 | 56.5 ± 3.7 | 76.3 ± 2.1 | 28.5 ± 2.5 | 45.8 ± 3.1 | SVM | 400 | 168 |
| CLIP-RN50x64 | 60.8 ± 2.6 | 70.2 ± 2.6 | 36.3 ± 4.5 | 48.8 ± 3.3 | SVM | 400 | 421 |
| CLIP-ViT-B32 | 60.4 ± 3.2 | 56.3 ± 3.5 | 33.1 ± 4.2 | 34.0 ± 3.6 | SVM | 400 | 86 |
| CLIP-ViT-I14 | 59.8 ± 2.1 | 69.4 ± 3.1 | 37.3 ± 3.7 | 40.2 ± 3.0 | SVM | 400 | 428 |
| MetaCLIP-b32-400m | 67.3 ± 2.6 | 68.1 ± 3.7 | 34.6 ± 2.8 | 43.3 ± 3.2 | SVM | 400 | 86 |
| MetaCLIP-b32-2.5b | 71.5 ± 3.0 | 72.5 ± 2.6 | 36.7 ± 3.2 | 40.4 ± 3.2 | SVM | 2500 | 86 |
| MetaCLIP-I14-400m | 62.9 ± 2.6 | 78.1 ± 3.5 | 37.1 ± 3.1 | 44.6 ± 4.0 | SVM | 400 | 304 |
| MetaCLIP-I14-2.5b | 80.8 ± 1.6 | 78.8 ± 1.8 | 40.2 ± 2.1 | 46.0 ± 1.5 | SVM | 2500 | 304 |
| MetaCLIP-h14-2.5b | 72.9 ± 3.5 | 78.9 ± 4.0 | 37.3 ± 1.5 | 43.1 ± 3.1 | SVM | 2500 | 632 |
| CLIP-RN101 | 37.5 | 52.1 | 22.9 | 22.9 | Zero-shot | 400 | 56 |
| CLIP-RN50 | 58.3 | 56.3 | 31.3 | 22.9 | Zero-shot | 400 | 38 |
| CLIP-RN50x4 | 62.5 | 54.2 | 25.0 | 33.3 | Zero-shot | 400 | 86 |
| CLIP-RN50x16 | 64.6 | 72.9 | 41.7 | 41.7 | Zero-shot | 400 | 168 |
| CLIP-RN50x64 | 66.7 | 70.8 | 37.5 | 47.9 | Zero-shot | 400 | 421 |
| CLIP-ViT-B32 | 64.6 | 54.2 | 39.6 | 18.8 | Zero-shot | 400 | 86 |
| CLIP-ViT-I14 | 79.2 | 62.5 | 37.5 | 35.4 | Zero-shot | 400 | 428 |
| MetaCLIP-b32-400m | 79.2 | 62.5 | 41.7 | 27.1 | Zero-shot | 400 | 86 |
| MetaCLIP-b32-2.5b | 83.3 | 75.0 | 35.4 | 35.4 | Zero-shot | 2500 | 86 |
| MetaCLIP-I14-400m | 83.3 | 75.0 | 43.8 | 41.7 | Zero-shot | 400 | 304 |
| MetaCLIP-I14-2.5b | 89.6 | 79.2 | 45.8 | 50.0 | Zero-shot | 2500 | 304 |
| MetaCLIP-h14-2.5b | 89.6 | 72.9 | 35.4 | 39.6 | Zero-shot | 2500 | 632 |

**Supplementary Fig. 3.**
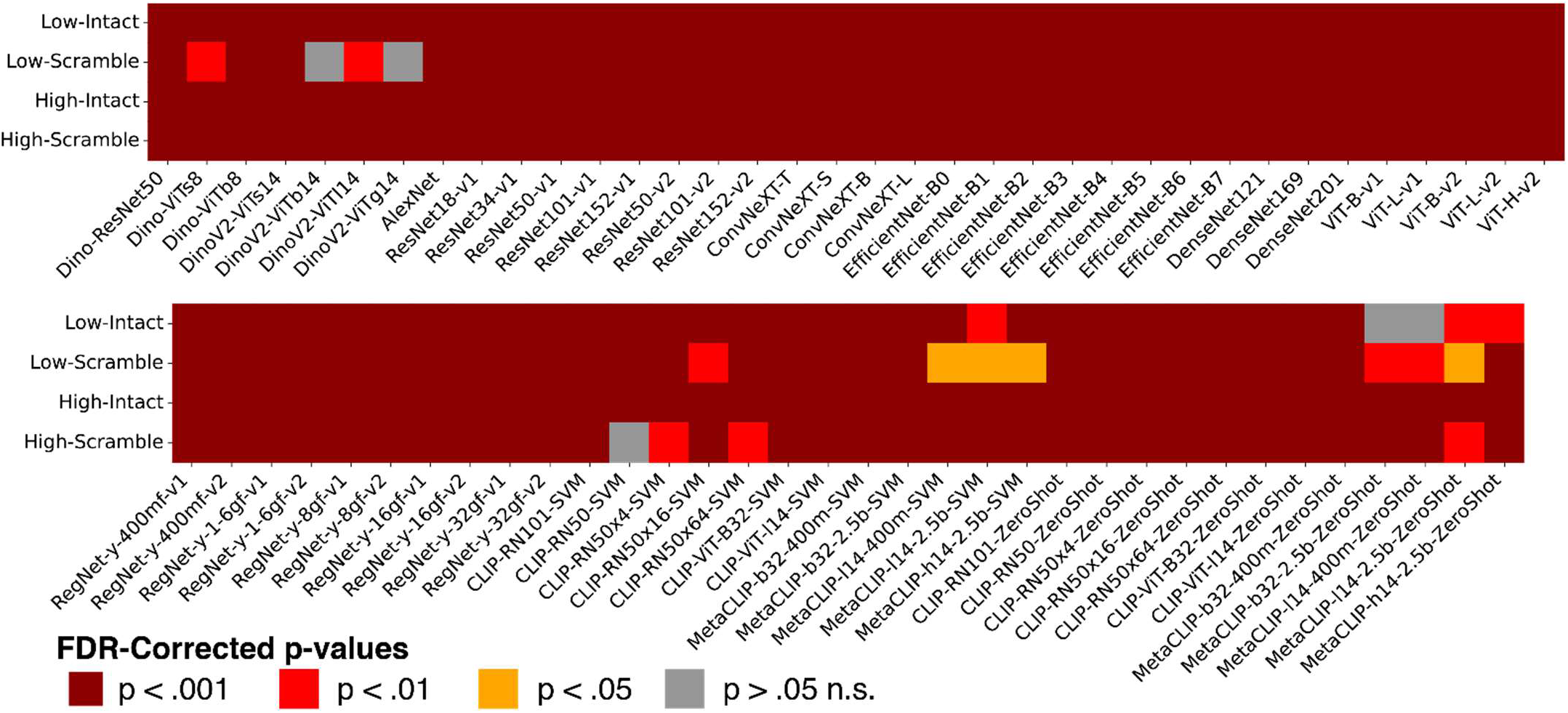
Heatmap of statistical comparison between model and human accuracy across conditions. Each cell indicates the FDR-corrected p-value from a non-parametric test comparing model accuracy to human accuracy in one of four experimental conditions (Low-Intact, Low-Scrambled, High-Intact, High-Scrambled). The horizontal axis lists all models evaluated. Color indicates statistical significance level after FDR correction. For SVM-evaluated models, accuracy distributions are based on 15 train/test splits; for zero-shot models, a single accuracy value was compared against human accuracies across 16 participants.

**Supplementary Table 2.** Statistical comparison of accuracy for each model group against human behavior (Supplementary Fig. 1). P-values are reported for low-difficulty intact, low-difficulty scramble, high-difficulty intact, and high-difficulty scramble conditions. Comparisons to human behavior were performed using two-sided Mann–Whitney U tests. Architecture comparisons were also conducted using two-sided Mann–Whitney U tests, except for the “self-supervised ResNet” group, which included only a single model; in this case, a two-sided Wilcoxon signed-rank tests was used. Asterisks (***) indicate p < 0.001. Direction of effect is denoted by arrows: (↑) indicates convolutional > transformer; (↓) indicates transformer > convolutional. Note: Some models groups had small sizes (e.g., n = 5), which may reduce statistical power and impact the interpretation of p-values.

| Model Group | Low-Intact | Low-Scramble | High-Intact | High-Scramble |
| --- | --- | --- | --- | --- |
| **Self-Supervised ResNet** | *** | *** | *** | *** |
| **Self-Supervised ViT** | *** | 0.016 | *** | *** |
| ViT vs. CNN | 0.15 n.s. | ↑ 0.03 | 0.15 n.s. | ↑ 0.03 |
| **Supervised CNNs** | *** | *** | *** | *** |
| **Supervised ViT** | *** | *** | *** | *** |
| ViT vs. CNN | ↓ 0.03 | 0.1 n.s. | ↓ 0.06 | ↓ 0.002 |
| **Language-Aligned ResNet SVM** | *** | 0.004 | *** | 0.01 |
| **Language-Aligned ViT SVM** | *** | 0.003 | *** | *** |
| ViT vs. CNN | ↑ 0.048 | 0.6 n.s. | 0.15 n.s. | 0.1 n.s. |
| **Language-Aligned ResNet Zero-Shot** | *** | 0.002 | *** | 0.002 |
| **Language-Aligned ViT Zero-Shot** | 0.24 n.s. | *** | *** | *** |
| ViT vs. CNN | ↑ 0.01 | 0.14 n.s. | 0.1 n.s. | 0.8 n.s. |

**Supplementary Table 3.** Statistical comparison of context effect magnitude (Fig. 2H) for each model group against human behavior and a no-effect baseline (zero). P-values are reported for both low- and high-difficulty conditions. Comparisons to human behavior were performed using two-sided Mann–Whitney U tests. Comparisons to the zero baseline were assessed using two-sided Wilcoxon signed-rank tests. Architecture comparisons were also conducted using two-sided Mann–Whitney U tests, except for the “self-supervised ResNet” group, which included only a single model; in this case, a two-sided Wilcoxon signed-rank tests was used. Asterisks (***) indicate p < 0.001. Direction of effect is denoted by arrows: (↑) indicates convolutional > transformer; (↓) indicates transformer > convolutional.

| Model Group | vs. Human |  | vs. Zero |  | Convolutional vs. Transformer |  |
| --- | --- | --- | --- | --- | --- | --- |
|  | Low Difficulty | High Difficulty | Low Difficulty | High Difficulty | Low Difficulty | High Difficulty |
| **Self-Supervised ResNet** | *** | *** | - | - | ↑ 0.03 | ↑ 0.03 |
| **Self-Supervised ViT** | *** | *** | 0.03 | 0.03 |  |  |
| **Supervised CNNs** | *** | *** | *** | 0.3 n.s. | 0.6 n.s. | ↓ 0.04 |
| **Supervised ViT** | *** | 0.004 | 0.06 | 0.12 n.s. |  |  |
| **Language-Aligned ResNet SVM** | 0.002 | *** | 0.06 | 0.06 | ↓ 0.047 | ↓ 0.018 |
| **Language-Aligned ViT SVM** | 0.14 n.s. | *** | 0.3 n.s. | 0.015 |  |  |
| **Language-Aligned ResNet Zero-Shot** | 0.29 n.s. | 0.003 | 0.63 n.s. | 0.59 n.s. | ↓ 0.006 | 0.2 n.s. |
| **Language-Aligned ViT Zero-Shot** | 0.004 | 0.017 | 0.016 | 0.46 n.s. |  |  |

## References

1. Aminoff, E. M., & Tarr, M. J. (2015). Associative Processing Is Inherent in Scene Perception. PLoS One, 10(6), e0128840. 10.1371/journal.pone.0128840

2. Bar, M. (2004). Visual objects in context. Nat Rev Neurosci, 5(8), 617–629. 10.1038/nrn1476

3. Biederman, I. (1972). Perceiving real-world scenes. Science, 177(4043), 77–80. 10.1126/science.177.4043.77

4. Brainard, D. H. (1997). The Psychophysics Toolbox. Spatial vision, 10 *4*, 433–436.

5. Brandman, T., & Peelen, M. V. (2017). Interaction between scene and object processing revealed by human fMRI and MEG decoding. Journal of Neuroscience, 37(32), 7700–7710.

6. Caron, M., Touvron, H., Misra, I., J’egou, H. e., Mairal, J., Bojanowski, P., & Joulin, A. (2021). Emerging Properties in Self-Supervised Vision Transformers. 2021 IEEE/CVF International Conference on Computer Vision (ICCV), 9630–9640.

7. Chen, L., Cichy, R. M., & Kaiser, D. (2022). Semantic Scene-Object Consistency Modulates N300/400 EEG Components, but Does Not Automatically Facilitate Object Representations. Cereb Cortex, 32(16), 3553–3567. 10.1093/cercor/bhab433

8. Cichy, R. M., Khosla, A., Pantazis, D., Torralba, A., & Oliva, A. (2016). Comparison of deep neural networks to spatio-temporal cortical dynamics of human visual object recognition reveals hierarchical correspondence. Scientific Reports, 6(1), 27755.

9. Cichy, R. M., Roig, G., Andonian, A., Dwivedi, K., Lahner, B., Lascelles, A., Mohsenzadeh, Y., Ramakrishnan, K., & Oliva, A. (2019). The Algonauts Project: A Platform for Communication between the Sciences of Biological and Artificial Intelligence. *ArXiv*, *abs/1905.05675*.

10. Collins, J. A., & Olson, I. R. (2014). Knowledge is power: how conceptual knowledge transforms visual cognition. Psychon Bull Rev, 21(4), 843–860. 10.3758/s13423-013-0564-3

11. Conwell, C., McMahon, E., Jagadeesh, A., Vinken, K., Sharma, S., Prince, J. S., Alvarez, G. A., Konkle, T., Livingstone, M., & Isik, L. (2025). Monkey See, Model Knew: Large Language Models Accurately Predict Visual Brain Responses in Humans and Non-Human Primates. BioRxiv, 2025.2003.2005.641284. 10.1101/2025.03.05.641284

12. Conwell, C., Prince, J. S., Kay, K. N., Alvarez, G. A., & Konkle, T. (2024). A large-scale examination of inductive biases shaping high-level visual representation in brains and machines. Nature Communications, 15(1), 9383.

13. Davenport, J. L., & Potter, M. C. (2004). Scene consistency in object and background perception. Psychol Sci, 15(8), 559–564. 10.1111/j.0956-7976.2004.00719.x

14. Doerig, A., Kietzmann, T. C., Allen, E. J., Wu, Y., Naselaris, T., Kay, K. N., & Charest, I. (2024). Visual representations in the human brain are aligned with large language models. arXiv preprint arXiv:2209.11737,

15. Dosovitskiy, A., Beyer, L., Kolesnikov, A., Weissenborn, D., Zhai, X., Unterthiner, T., Dehghani, M., Minderer, M., Heigold, G., Gelly, S., Uszkoreit, J., & Houlsby, N. (2020). An Image is Worth 16×16 Words: Transformers for Image Recognition at Scale. ArXiv, abs/2010.11929.

16. Gauthier, I., James, T. W., Curby, K. M., & Tarr, M. J. (2003). The influence of conceptual knowledge on visual discrimination. Cogn Neuropsychol, 20(3), 507–523. 10.1080/02643290244000275

17. Ge, Y., Tang, Y., Xu, J., Gokmen, C., Li, C., Ai, W., Martinez, B. J., Aydin, A., Anvari, M., & Chakravarthy, A. K. (2024). BEHAVIOR Vision Suite: Customizable Dataset Generation via Simulation. Proceedings of the IEEE/CVF Conference on Computer Vision and Pattern Recognition,

18. Geirhos, R., Narayanappa, K., Mitzkus, B., Bethge, M., Wichmann, F., & Brendel, W. (2020). On the surprising similarities between supervised and self-supervised models. ArXiv, abs/2010.08377.

19. Gilbert, C. D., & Li, W. (2013). Top-down influences on visual processing. Nature Reviews Neuroscience, 14(5), 350–363. 10.1038/nrn3476

20. He, K., Zhang, X., Ren, S., & Sun, J. (2015). Deep Residual Learning for Image Recognition. 2016 IEEE Conference on Computer Vision and Pattern Recognition (CVPR), 770–778.

21. Hebart, M. (2009). Stimulus Phase Scrambling Code. http://martin-hebart.de/webpages/code/stimuli.html

22. Hebart, M. N., Contier, O., Teichmann, L., Rockter, A. H., Zheng, C. Y., Kidder, A., Corriveau, A., Vaziri-Pashkam, M., & Baker, C. I. (2023). THINGS-data, a multimodal collection of large-scale datasets for investigating object representations in human brain and behavior. Elife, 12. 10.7554/eLife.82580

23. Jacob, G., Pramod, R. T., Katti, H., & Arun, S. P. (2021). Qualitative similarities and differences in visual object representations between brains and deep networks. Nature Communications, 12(1), 1872. 10.1038/s41467-021-22078-3

24. Kaiser, D., Quek, G. L., Cichy, R. M., & Peelen, M. V. (2019). Object vision in a structured world. Trends in cognitive sciences, 23(8), 672–685.

25. Khaligh-Razavi, S.-M., & Kriegeskorte, N. (2014). Deep supervised, but not unsupervised, models may explain IT cortical representation. PLoS computational biology, 10(11), e1003915.

26. Kreiman, G., & Serre, T. (2020). Beyond the feedforward sweep: feedback computations in the visual cortex. Ann N Y Acad Sci, 1464(1), 222–241. 10.1111/nyas.14320

27. Krizhevsky, A., Sutskever, I., & Hinton, G. E. (2012). ImageNet classification with deep convolutional neural networks. Communications of the ACM, 60, 84–90.

28. Li, C., Zhang, R., Wong, J., Gokmen, C., Srivastava, S., Martfn-Martfn, R., Wang, C., Levine, G., Lingelbach, M., & Sun, J. (2023). Behavior-1k: A benchmark for embodied ai with 1,000 everyday activities and realistic simulation. Conference on Robot Learning,

29. Lupyan, G., Abdel Rahman, R., Boroditsky, L., & Clark, A. (2020). Effects of Language on Visual Perception. Trends in cognitive sciences, 24(11), 930–944. 10.1016/j.tics.2020.08.005

30. Lupyan, G., Thompson-Schill, S. L., & Swingley, D. (2010). Conceptual penetration of visual processing. Psychol Sci, 21(5), 682–691. 10.1177/0956797610366099

31. Malcolm, G. L., Groen, I. I. A., & Baker, C. I. (2016). Making Sense of Real-World Scenes. Trends Cogn Sci, 20(11), 843–856. 10.1016/j.tics.2016.09.003

32. Markov, Y. A., & Võ, M. L.-H. (2025). Scene consistency enhances state representations of real-world objects. Scientific Reports, 15(1), 18581. 10.1038/s41598-025-01662-3

33. Munneke, J., Brentari, V., & Peelen, M. V. (2013). The influence of scene context on object recognition is independent of attentional focus. Front Psychol, 4, 552. 10.3389/fpsyg.2013.00552

34. Oliva, A., & Torralba, A. (2007). The role of context in object recognition. Trends Cogn Sci, 11(12), 520–527. 10.1016/j.tics.2007.09.009

35. Oquab, M., Darcet, T., Moutakanni, T., Vo, H. Q., Szafraniec, M., Khalidov, V., Fernandez, P., Haziza, D., Massa, F., El-Nouby, A., Assran, M., Ballas, N., Galuba, W., Howes, R., Huang, P.-Y., Li, S.-W., Misra, I., Rabbat, M. G., Sharma, V.,…Bojanowski, P. (2023). DINOv2: Learning Robust Visual Features without Supervision. ArXiv, abs/2304.07193.

36. Peelen, M. V., Berlot, E., & de Lange, F. P. (2024). Predictive processing of scenes and objects. Nat Rev Psychol, 3, 13–26. 10.1038/s44159-023-00254-0

37. Peters, B., & Kriegeskorte, N. (2021). Capturing the objects of vision with neural networks. Nature Human Behaviour, 5(9), 1127–1144. 10.1038/s41562-021-01194-6

38. Radford, A., Kim, J. W., Hallacy, C., Ramesh, A., Goh, G., Agarwal, S., Sastry, G., Askell, A., Mishkin, P., Clark, J., Krueger, G., & Sutskever, I. (2021). Learning Transferable Visual Models From Natural Language Supervision. International Conference on Machine Learning,

39. Rajaei, K., Mohsenzadeh, Y., Ebrahimpour, R., & Khaligh-Razavi, S. M. (2019). Beyond core object recognition: Recurrent processes account for object recognition under occlusion. PLoS Comput Biol, 15(5), e1007001. 10.1371/journal.pcbi.1007001

40. Schrimpf, M., Kubilius, J., Hong, H., Majaj, N. J., Rajalingham, R., Issa, E. B., Kar, K., Bashivan, P., Prescott-Roy, J., Schmidt, K., Yamins, D., & DiCarlo, J. J. (2018). Brain-Score: Which Artificial Neural Network for Object Recognition is most Brain-Like? BioRxiv.

41. Ullman, S., Assif, L., Fetaya, E., & Harari, D. (2016). Atoms of recognition in human and computer vision. Proc Natl Acad Sci U S A, 113(10), 2744–2749. 10.1073/pnas.1513198113

42. Võ, M. L. (2021). The meaning and structure of scenes. Vision Res, 181, 10–20. 10.1016/j.visres.2020.11.003

43. Võ, M. L., & Wolfe, J. M. (2013). Differential electrophysiological signatures of semantic and syntactic scene processing. Psychol Sci, 24(9), 1816–1823. 10.1177/0956797613476955

44. Wang, A. Y., Kay, K., Naselaris, T., Tarr, M. J., & Wehbe, L. (2023). Better models of human high-level visual cortex emerge from natural language supervision with a large and diverse dataset. Nature Machine Intelligence, 5(12), 1415–1426.

45. Wilcoxon, F. (1945). Individual Comparisons by Ranking Methods. Biometrics, 1, 196–202.

46. Wischnewski, M., & Peelen, M. V. (2021). Causal neural mechanisms of context-based object recognition. Elife, 10, e69736.

47. Xu, H., Xie, S., Tan, X. E., Huang, P.-Y., Howes, R., Sharma, V., Li, S.-W., Ghosh, G., Zettlemoyer, L. S., & Feichtenhofer, C. (2023). Demystifying CLIP Data. *ArXiv*, *abs/2309.16671*.

48. Yamins, D. L., Hong, H., Cadieu, C. F., Solomon, E. A., Seibert, D., & DiCarlo, J. J. (2014). Performance-optimized hierarchical models predict neural responses in higher visual cortex. Proc Natl Acad Sci U S A, 111(23), 8619–8624. 10.1073/pnas.1403112111

49. Zhou, B., Bau, D., Oliva, A., & Torralba, A. (2019). Interpreting Deep Visual Representations via Network Dissection. IEEE Transactions on Pattern Analysis and Machine Intelligence, 41(9), 2131–2145. 10.1109/TPAMI.2018.2858759

50. Zhuang, C., Xiang, V., Bai, Y., Jia, X., Turk-Browne, N., Norman, K., DiCarlo, J. J., & Yamins, D. L. K. (2022). How Well Do Unsupervised Learning Algorithms Model Human Real-time and Life-long Learning? Adv Neural Inf Process Syst, 35, 22628–22642.

